# Insights on the historical biogeography of Philippine native pigs and its relationship with Continental domestic and wild boars

**DOI:** 10.1101/2021.07.23.453525

**Authors:** John King N. Layos, Ronel N. Geromo, Dinah M. Espina, Masahide Nishibori

## Abstract

The Philippine archipelago was believed to have never been connected to the Asian continent even during the severe Quaternary sea-level drops. As a result, the history of pig dispersal in the Philippines remains controversial and must have some anthropogenic origin associated with some human migration events. In this study, the context of origin, dispersal, and the level of genetic introgression in Philippine native pigs were deduced using mitochondrial DNA D-loop analysis altogether with domestic pigs and wild boars corresponding to their geographic origin. Results revealed a considerable genetic diversity (0.900±0.016), and a widespread Asian pig-ancestry (94.60%) were revealed in the phylogenetic analysis with admixed European pig-origin (5.10%) harboring various fractions of ancestry from Berkshire and Landrace. The close genetic connection between the continental wild boars and domestic pigs present in the Philippine pigs corroborates our hypothesis of a genetic signal that could potentially be associated with the recently reported multiple waves of human migrations to the Philippines during the last 50,000 years. The high frequency of haplotypes (54.08%) that collapsed in the D7 haplogroup represent an interesting challenge as its distribution does not coincide with the hypothesized migratory route of the Neolithic Austronesian-speaking populations. We detected the first Pacific Clade signature and ubiquitously distributed D2 haplotypes which postulate the legitimate dispersal of pigs associated with the multiple waves of human migrations involving the Philippines. The multimodal mismatch and neutrality test statistics both Fu’s F*s* and Tajima’s D correlates the long stationary period of effective population size revealed in the Bayesian skyline plot. While the sudden decrease in population was consistent with the pronounced population bottleneck of Asian and European pigs during the interglacial periods of the Pleistocene.

## Introduction

The wild boar (*Sus scrofa* L.), recognized as the ancestor of the domestic pig, is one of the most widely distributed mammals found throughout Eurasia, including South and East Asia, and extending to North Africa. This species was also introduced into the Americas, Australia and Oceania [1]. Because of its relationship with human settlement and movement, studies on the phylogeography of *S. scrofa* have provided significant evidence revealing both anthropological and biogeographical history [2]. From the point of view of molecular phylogeny at a larger geographical scale, wild boars are genetically divided into Asian and European clades [3-6], which have split during the Mid-Pleistocene 1.6–0.8 Ma ago [7]. Wild boars from East and South-Eastern Asia predominantly have greater amounts of genetic variation than European wild boars, based on both mtDNA [4] and nuclear markers [8]. Island South-Eastern Asia (ISEA) and mainland South-Eastern Asia (MSEA), known to be the area of phylogenetic origin of wild boars, is a biodiversity hotspot where most other species in the genus *Sus* are present [9].

The Philippines is one of the most biologically rich regions in the world with exceptionally high levels of endemism for a country of its size. It has repeatedly been tagged as a region of global conservation priority – a top hotspot for both terrestrial and marine ecosystems [10-13]. The Philippine archipelago was believed to have never been connected to the Asian continent during the past glacial periods [14] hence, *S. scrofa* was unable to reach the archipelago from the MSEA [15-16]. Therefore, the *S. scrofa* that exists in the Philippines today might suggest some anthropogenic origin, likely arriving through human migration events [17]. The Austronesian settlers first colonized the Philippine archipelago around 4,000 years ago [18-19] and were believed to have initiated the dispersal and translocation of pigs in the country. From the Philippines they dispersed fairly rapidly to south and west into the ISEA [18-19]. They also moved east into the Marianas, and the Bismarck Archipelago, where the Lapita Complex developed around ca. 3300-3150 cal. BP. From there, they travelled further east into the Solomon Islands and Vanuatu, and eventually spread further into Remote Oceania [19]. However, the circumstantial lack of corroborating archaeological evidence supporting the introduction of pigs in the Philippines have long been daunting. Numerous archaeologists and geneticists have argued that connections with MSEA, as opposed to the “Out of Taiwan” model of dispersal are responsible for the introduction of the Austronesian languages and agriculture in ISEA [20-21]. This overlapping hypothesis was further aggravated by the absence of Pacific clade signatures both in Taiwan and the Philippines. For this reason, [22] precluded the Philippines as the point of departure further eastwards into the Pacific for domestic pigs. These haplotypes appear to have originated somewhere in peninsular Southeast Asia and were transported through Malaysia, Sumatra, Java, and islands in Wallacea such as Flores, Timor, and the Moluccas [4,22-24], and have shown strong support for the connection of Neolithic material culture between Vietnam and ISEA [20]. However, in contrast, some modern and ancient Philippine pigs possessed a unique haplotype stemming from the island of Lanyu (Orchid Island), located between the northern Philippines and southern Taiwan [25]. Despite these complex genetic evidence, molecular studies undertaken on pigs in the Philippines are still very limited thus, the pattern of pig expansion and dispersal remains ambiguous. Therefore, modern pig genetic studies could help elucidate the ongoing discussion on pig dispersal and migration involving the Philippines. Hence, this study aims at describing the mtDNA variability, genetic structure and phylogeographic origin of Philippine pigs, understanding if the currently observed genetic diversity and structure have a signature of past demographic expansion in reference to those observed in the MSEA and finally, contribute relevant insights to the conflicting hypothesis proposed to explain the introduction of pigs involving the Philippines.

## Materials and Methods

### Sampling and laboratory analysis

In this study, blood samples and hair follicles of Philippine native pigs and wild pigs were sampled [S1 Appendix] in accordance with institutional, local and national guidelines regarding animal care and use in experimentation established by the Laboratory of Animal Genetics, Hiroshima University (No. 015A170426). Samples were preserved in tubes kept in - 20°C. Photographs were obtained to document the morphological characteristics and differences within these pig populations (Fig 1). The genomic DNA was extracted using the phenol-chloroform method following the recommended protocol described by [26]. The 5.0-kbp mitochondrial DNA (mtDNA) fragment were amplified using a long and accurate – PCR kit (KOD FX-neo Polymerase, TOYOBO, Otsu, Japan) using the established primer set: Sus mt. 5.0 FL-2: 5’-ATGAAAAATCATCGTTGTACTTCAACTACAAGAAC-3’; Mum R: 5’-TTCAGACCGACCGGAGCAATCCAGGTCGGTTTCTATCTA-3’. The reaction began with an initial denaturation at 94°C for 2 min, followed by 30 cycles of denaturation at 98°C for 10 sec, annealing at gradients at 57°C for 30 sec, and primer extension at 68°C for 2 min and 30 sec. The last step was 8 min final extension period at 68°C. For mtDNA displacement (D-loop) region amplification, the ca. 1.3 kbp fragment was amplified using another primer set, Sus mtD F1: AACTCCACCATCAGCACCCAAAG, Sus mtD R1: CATTTTCAGTGCCTTGCTTTGATA [27]. The reaction began with an initial denaturation at 94°C for 2 min, after that, followed by 30 cycles of denaturation at 98°C for 10 sec, annealing at gradients 59°C for 30 sec, and extension at 68°C for 30 sec. The last step was a 5 min final extension period at 68°C. The amplification was done using the GeneAmp PCR System 9700 (Applied Biosystems, Foster City, CA, USA). The PCR products from the segmental amplification were cleaned and purified using Exonuclease I (ExoI) and Shrimp Alkaline Phosphatase (SAP) to degrade the residual PCR primers and dephosphorylate the remaining dNTPs, respectively. After this, samples were sequenced using ABI3130 sequencer for direct DNA sequencing and fragment analysis.

**Fig 1.**
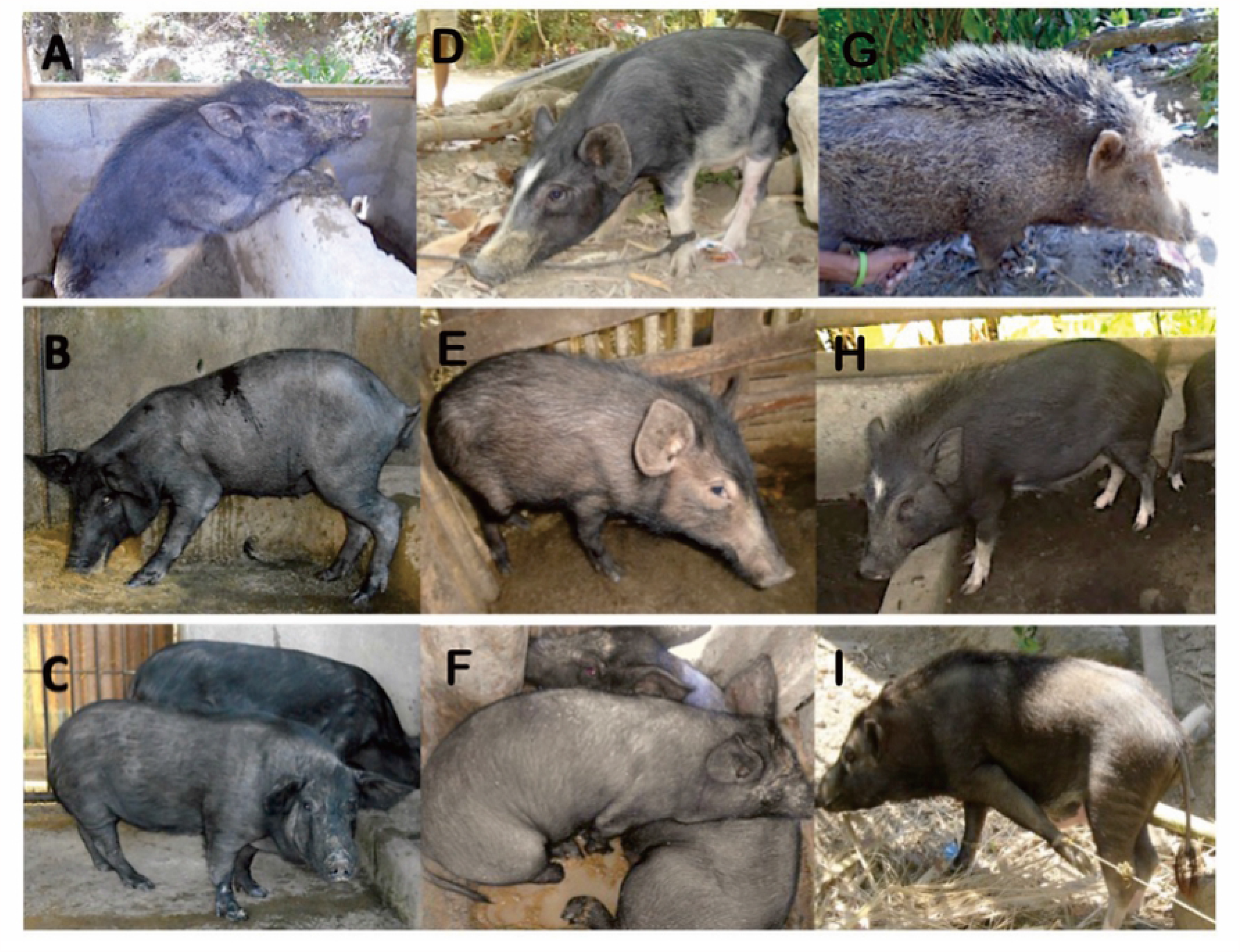
A sample of the morphological variations across Philippine native pigs. a) Mambusao, Capiz. b) Barbaza, Antique. c) Dao, Capiz. d) Buenavista, Guimaras. e) Dingle, Iloilo. f) Nueva Valencia, Guimaras. g) Bugasong, Antique. h) Sebaste, Antique. i) Barbaza, Antique.

### Phylogenetic and population structure analysis

The profile alignments of the mtDNA D-loop sequenced data were done through CLUSTAL W algorithm as implemented in the Molecular Evolutionary Genetics Analysis (MEGA) [28]. About 1044 bp of the control region sequences were aligned and edited until one highly variable tandem repeat motif (5’-CGTGCGTACA-3’) remained. Haplotype sequences were submitted to GenBank National Center for Biotechnology Information (NCBI) databases (MN625805-MN625830; MW924902-MW92973). To place our results in a broader context, sequences obtained from this study were shortened from the original size of 1044 bp to 510 bp to allow for comparison of the pooled sequences from the previously documented sequences from MSEA available in GenBank [S2 Appendix]. The diversity measures such as the number of polymorphic segregating sites (S), haplotype diversity (hd) and nucleotide diversity (π) were estimated by DNA Sequence Polymorphism (DnaSP) 5.10 software [29]. The genetic structure of the studied populations was assessed employing the analysis of molecular variance (AMOVA) as implemented in ARLEQUIN. Four genetic structure hypotheses based on geographical locations were tested namely, (1) Philippine pigs (no groupings); (2) Philippines vs. MSEA combined; (3) Philippines *vs*. Bhutan *vs*. Myanmar, Laos, Cambodia, and Vietnam; (4) Philippines *vs*. Bhutan and Myanmar *vs*. Cambodia, Lao and Vietnam. The extent of population genetic differentiation was further quantified by F_*ST*_ statistic using ARLEQUIN [30]. Fixation index (Φ) statistics of population genetics were calculated, and the significance of the variance component were performed using 1,000 random permutations. Φ_CT_ is the difference among the groups of total haplotypes, Φ_SC_ is the difference among populations within groups, and Φ_ST_ is the difference among localities within populations. The AMOVA estimate genetic structure indices were determined using information on the allelic content of haplotypes as well as their frequencies [31]. All haplotypes were combined including the downloaded sequences represented animals classified as domestic and wild *S. scrofa* from Europe and Asia [S3 Appendix] for phylogeny reconstruction using the Maximum Likelihood (ML) inference with GTR+G+I as the best fitted model using PhyML v.3.0 [32] as Warthog (*Phacochoerus africanus*; DQ409327) as an outgroup. Bootstrap values were estimated with 1,000 repetitions. To provide a more detailed information on the phylogenetic relationship among these haplotypes, a reduced median network was constructed using Network v.4.1 [33-34], available at http://www.fluxus-engineering.com. This method calculates the net divergence of each taxon from all other taxa as the sum of the individual distances from variance within and among groups of Philippine pigs and comparison sequences. The nomenclatures described by Larson et al. (2005) with six clades (D1 to D6) including the newly proposed clade by Tanaka et al. (2008) were used as the reference for the clade notation.

### Population demographic analysis

Demographic history was inferred by the analysis of the distribution of the number of site differences between pairs of sequences (mismatch distribution), which was carried out on the previously described pooled samples, as implemented in DnaSP 5.10 software [29]. Expected values for a model of population growth-decline were calculated and plotted against the observed values. Populations that have experienced a rapid demographic growth in the past show unimodal distributions, whereas those at demographic equilibrium or decline presents multimodal distributions [35]. Harpendings [36] raggedness index (H*ri*; quantifying the smoothness of the mismatch distributions and distinguishing between population expansion and stability) and the sum of squared deviations (SSD) (1,000 simulated samples of pairwise nucleotide differences), as implemented in ARLEQUIN [31], were used to evaluate the Rogers [37] sudden expansion model, which fits to a unimodal mismatch distribution [35]. To test for population expansion, we employed three other tests: Fu’s [38] F_*S*_ and Tajima’s D statistical tests using ARLEQUIN and testing their significance over 1,000 permutations; and Ramos-Onsins and Rozas R^2^ test [39] by means of DnaSP. Statistical tests and confidence intervals for F_*S*_ were based on a coalescent simulation algorithm and for R^2^ on parametric bootstrapping with coalescence simulations.

The past population dynamics were also explored with the Bayesian skyline plot (BSP) [40] model with standard Markov chain Monte Carlo sampling procedure (MCMC) under HKY + G model of substitution [41] with four gamma categories using Beast v.2.6.0 [42]. The BSP represents population size changes over time, inferred with mtDNA and the assumed mutation rate. Two independent analyses were performed using all sequences from this study and the 130 sequences from MSEA using 1.36 × 10^−8^ mutation rate (mutation rate per nucleotide site per year according to previously estimates for mammalian mtDNA control region; [43]) under the strict clock. The MCMC analysis was run for 50,000,000 generations. Independent runs (logs and trees) were pooled using Log Combiner, discarding burn-in of the first 10% and parameter values were sampled every 5,000 generations. Tracer v.1.7 as used to confirm correct MCMC chain convergence with an effective sample size (ESS) > 200 within the Log files, and to visualize the dynamics of the effective population size over time. The light-blue shaded area marks the 95% highest posterior density (HPD). The X-axes are time in thousands of years before present (BP) and the Y-axes are the mean effective population size in millions of individuals divided by generation time on a log scale.

## Results

### Genetic diversity and population differentiation

Overall, 236 sequences including the 106 Philippine native pigs (two wild pigs were excluded) were used to measure population genetic structure and differentiation. The nucleotide sequences were aligned relative to the representative haplotypes of Asian domestic pigs under accession number AB041480 [S4 Appendix]. In the alignment of sequences, all the variable sites represented substitution mutation. Overall, 23 haplotypes were detected from the Philippines (PHL), 11 in Cambodia, 9 in Bhutan, 10 in Laos, 17 in Myanmar, and 4 in Vietnam (Table 1). These sequences collapsed when pooled from 76 to 57 haplotypes. Twenty-six (6 PH and 20 MSEA) of the 57 haplotypes were represented by a single sequence. The highest number of shared individuals was noted in PH37 haplotype consisting of 31 sequences (25 PH and 6 MSEA; 13.59%) which corresponds to D7 haplogroup and was previously referred as the mitochondrial Southeast Asian haplogroup (MTSEA) [44]. The haplotype diversity of native and domestic pigs in MSEA when combined was 0.966±0.006 with Myanmar showing the highest haplotype diversity (0.958±0.018), followed by Laos with 0.925±0.047 and the Philippines with 0.900±0.016. Meanwhile, the lowest haplotype diversity was observed from Vietnam (0.800±0.172). On the other hand, nucleotide diversity (π) was highest in the Philippines (0.012±0.006), closely similar to Bhutan (0.010±0.006), while the lowest was noted in Vietnam pigs (0.004±0.003). However, the small sample number size of pigs from Vietnam may account for its apparent low genetic diversity.

**Table 1.** mtDNA indices of Philippine native pigs and mainland Southeast Asian pigs

| COUNTRIES | n | vs | P | S | #h | MPD | h | $\pi$ |
| --- | --- | --- | --- | --- | --- | --- | --- | --- |
| KHM | 49 | 13 | 9 | 4 | 11 | $4.272 \pm 2.154$ | $0.854 \pm 0.028$ | $0.008 \pm 0.005$ |
| BTN | 30 | 16 | 9 | 7 | 9 | $4.855 \pm 2.436$ | $0.807 \pm 0.051$ | $0.010 \pm 0.006$ |
| LAO | 16 | 12 | 8 | 4 | 10 | $3.867 \pm 2.049$ | $0.925 \pm 0.047$ | $0.008 \pm 0.005$ |
| MMR | 29 | 18 | 11 | 7 | 17 | $4.502 \pm 2.282$ | $0.958 \pm 0.018$ | $0.009 \pm 0.005$ |
| VNM | 6 | 4 | 2 | 2 | 4 | $2.000 \pm 1.304$ | $0.800 \pm 0.172$ | $0.004 \pm 0.003$ |
| TOTAL | 130 | 37 | 23 | 14 | 44 | | $0.966 \pm 0.006$ | $0.010 \pm 0.006$ |
| PHL | 106 | 29 | 25 | 4 | 22 | $5.207 \pm 2.542$ | $0.900 \pm 0.016$ | $0.012 \pm 0.006$ |

PHL= Philippines; KHM= Cambodia; BTN= Bhutan; MMR= Myanmar; VNM= Vietnam; n= number of samples; vs= variable sites; P= parsimony informative sites; S= singleton variable sites; #h= number of haplotypes; MPD= mean pairwise differences (SD); h= gene diversity: haplotype level (SD); $\pi$ = nucleotide diversity (SD).

Based on the AMOVA of mtDNA D-loop data, all hypothesis subjected for analysis revealed a significant population subdivision (Φ_ST_ values, *p*<0.01) which suggests a distinct genetic structure in all studied geographical locations (Table 2). The inference of genetic differentiation was equally evident in both the neighbor-joining and network analyses (Figs 2 and 3). The pairwise F_*ST*_ estimate ranged from 0.050 to 0.476 (Table 3A) showed significantly higher genetic differentiation (Φ_ST_ values, *p*<0.01) observed between most pigs in the studied populations and between Philippine islands. However, insignificant and small pairwise Φ_ST_ estimates were noticed in Laos and Cambodian pigs (Φ_ST_=0.050; *p*=0.0631), indicating that pigs from these countries were not isolated from each other. The corrected average pairwise differences revealed consistent tendencies with the pairwise Φ_ST_ (Table 3B).

**Table 2.** Analysis of molecular variance (AMOVA) of Philippine native pigs and mainland Southeast Asian pigs

| HYPOTHESIS | Sources of variation | % Variation | $\Phi$ |
| --- | --- | --- | --- |
| PHL pigs<br>(No groupings) | Among populations | 16.95** | $\Phi_{ST} = 0.16952$ |
| | Within populations | 83.05 | $\Phi_{SC} = 0.83052$ |
| PHL vs. MSEA<br>combined | Among groups | -9.34 | $\Phi_{CT} = -0.09338$ |
| | Among populations within groups | 21.47** | $\Phi_{SC} = 0.19639$ |
| | Within populations | 87.87** | $\Phi_{ST} = 0.12135$ |
| PHL vs. BTN vs. MMR,<br>VNM, KHM, LAO | Among groups | 0.01 | $\Phi_{CT} = 0.00006$ |
| | Among populations within groups | 14.61** | $\Phi_{SC} = 0.14614$ |
| | Within populations | 85.38** | $\Phi_{ST} = 0.14619$ |
| PHL vs. BTN, MMR vs.<br>VNM, KHM, LAO | Among groups | 2.51 | $\Phi_{CT} = 0.14837$ |
| | Among populations within groups | 12.33** | $\Phi_{SC} = 0.12644$ |
| | Within populations | 85.16** | $\Phi_{ST} = 0.14837$ |

\*\* $p < 0.01$ as tested by randomization (1000 permutations) using ARLEQUIN; MSEA = Mainland Southeast Asia; PHL = Philippines; KHM = Cambodia; BTN = Bhutan; MMR = Myanmar; VNM = Vietnam; $\Phi_{CT}$ = the difference among the groups of total haplotypes; $\Phi_{SC}$ = the difference among populations within groups; $\Phi_{ST}$ = the difference among localities within populations.

**Fig 2.**
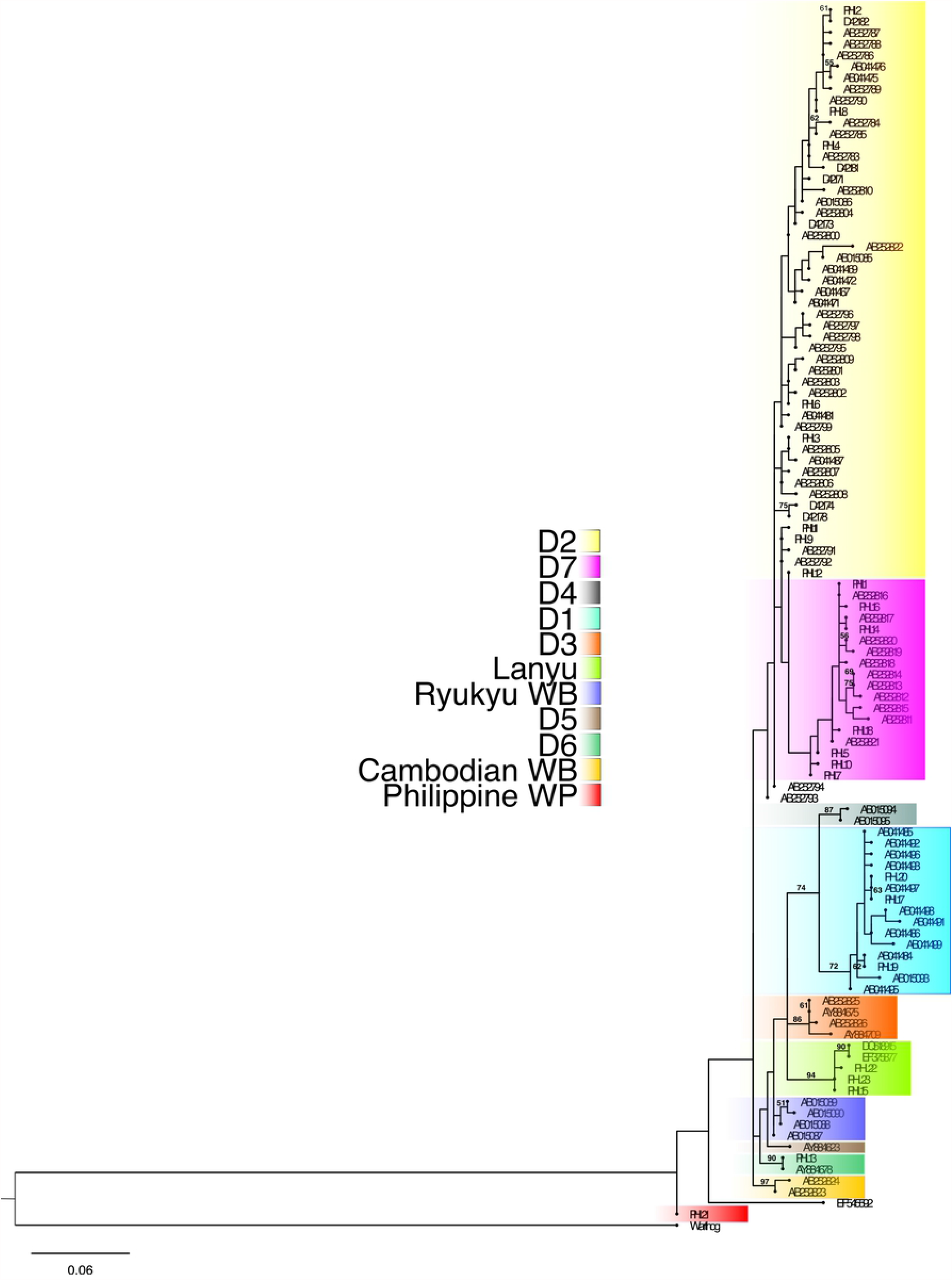
**Phylogenetic relationships of Philippine native pigs and wild pigs with continental domestic and wild boar. The number indicated in the nodes were bootstrap supports based on 1,000 replicates with warthog as the outgroup. Bootstrap values lower that 50% were not shown. Philippine pigs revealed to comprised founder sources from five different geographic origins excluding the endemic Philippine wild pigs.**

**Fig 3.**
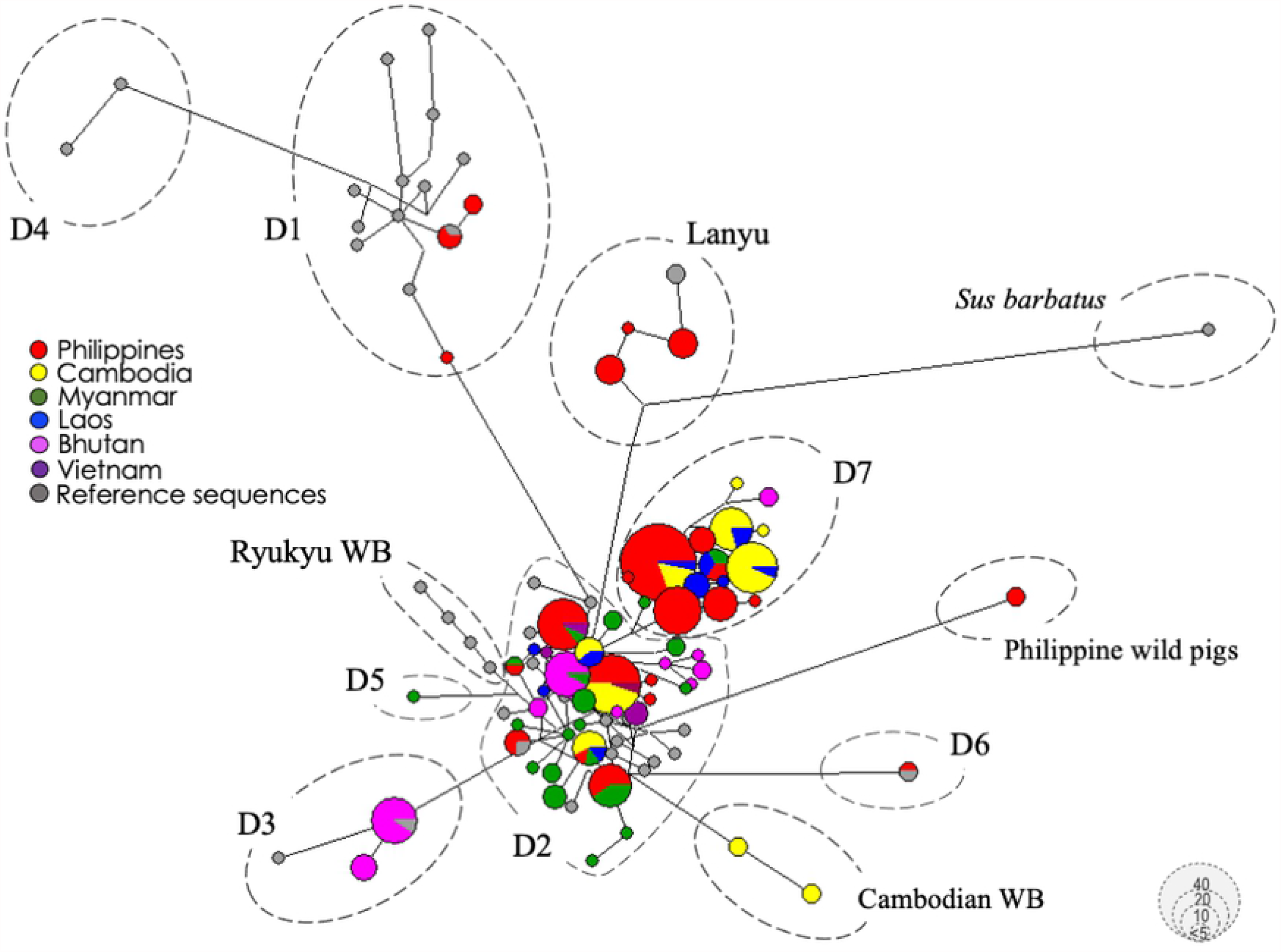
**The median-joining network of Asian and European pig haplotypes including the global reference sequences showing haplogroup classification. Some haplotypes clustered together coinciding with their geographic area of origin, while selected haplotypes diverse and shared by individuals of different breeds from different geographical regions, indicating a negative correspondence between the geographic origin and the relationships among breeds. The size of each circle is proportional to the haplotype frequency. Color represents regions of sequence origin.**

**Table 3.**
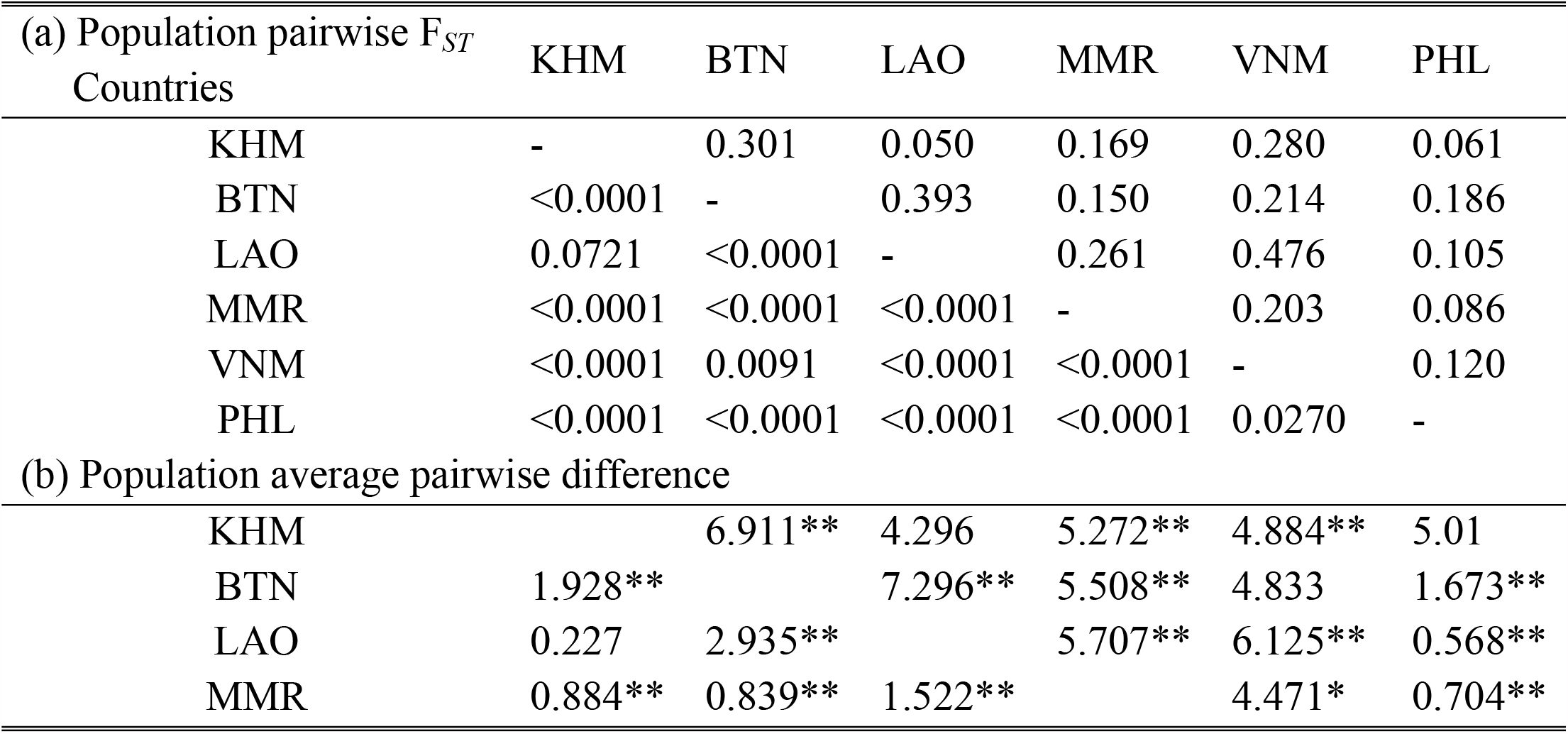
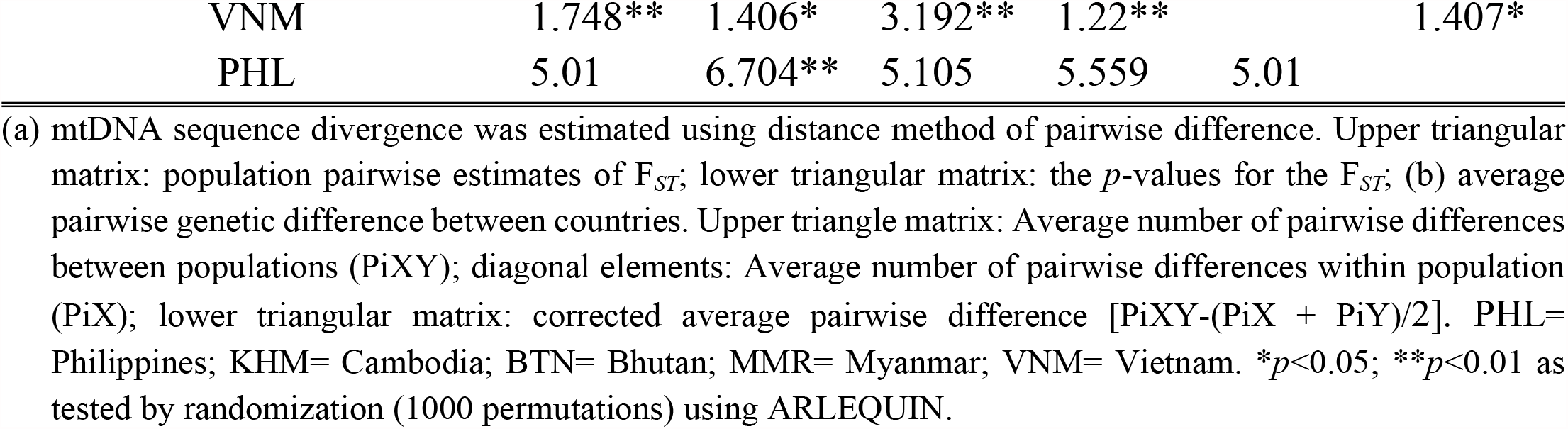
Genetic divergence among populations of Philippine native pigs and mainland Southeast Asian pigs

To investigate the relationship of Philippine native pigs from those Asian and European pigs, we included 40 sequences [4] to accommodate most of the major mtDNA porcine haplotypes. Overall, a total of 285 sequences were used to perform the phylogenetic tree and median-joining network analysis. The NJ tree branched into two core lineages, one of Asian phylogeographic origins and one of European phylogeographic origins (Fig 2). Network analysis generally supported the phylogenetic tree and revealed a strong genetic structuring among Philippine pig haplotypes where different phylogroups could be observed (Fig 3). A widespread Asian ancestry (94.90%) was observed in all Philippine pig haplotypes studied. Seven (PHL1, 5, 7, 8, 14, 16 and 18) of 21 Philippine pig haplotypes nested under the D7 haplogroup (referred earlier as MTSEA), which consists of 54.08% of the total studied population. This haplogroup was earlier reported in the MSEA as restricted to the Indo-Burma Biodiversity Hotspots (IBBH) [44] and as a distinct clade and which was not described in previous studies [4]. Eight haplotypes (PHL2, 3, 4, 6, 8, 9, 11 and 12; 36.73%) were distributed in the D2 haplogroup which corresponds to what [5] recognized as widely distributed Chinese domestic pigs, a global pig breed that has some relationship with Asian pigs, as well as with East Asian wild boars [45-46]. Three haplotypes (PHL17, 19 and 20; 5.10%) revealed the presence of admixed ancestry in D1 haplogroup (European clade), harboring different fractions of maternal lineages from Berkshire and Landrace. Intriguingly, one and three haplotypes clustered under the Pacific Clade (PHL13) and the distinct Type I Lanyu pig (PHL15, 22 and 23), respectively. As we shall discuss later, there had been no similar haplotypes reported in previous studies on the existence of Pacific clade haplotypes in the Philippines. One haplotype (PHL21) of a Philippine wild pig cannot be classified under any of the proposed haplogroups and formed a vague cluster from the nomenclature quotation of porcine mtDNA control region haplotypes found in the previous researches. It possessed a unique nucleotide polymorphism at sites T54C, C127T, A148G, T406C, G407A and a transversion substitution at the base G88T [S4 Appendix]. Thus, we propose that the mtDNA haplotypes in these individuals should be classified into a distinct cluster with a potential for recognizing novel subspecies of wild pigs in the Philippines.

### Past population dynamics

The mismatch distributions were also calculated to investigate the hypothesis of population expansion. The distribution of pairwise nucleotide differences of the studied Philippine pig populations revealed multimodal patterns of mismatch distribution (Fig 4A), which may suggest a population in decline or stable demographic equilibrium. On the contrary, pigs in MSEA combined revealed a unimodal mismatch distribution, significant and large negative Fu’s F_*S*_ value (−25.282; *p*<0.01), together with the small and non-significant value of Harpending’s raggedness index (H*ri*), likely supporting a scenario of demographic expansion experienced in the past (Fig 4B; Table 4). When the effects of natural selection and past demographic changes are examined in each of the populations per country using the Tajima’s D and Fu’s F_*S*_ estimates, only Myanmar had a significantly negative Fu’s F_*S*_ values (−6.542; *p*<0.05) while all Tajima’s D values for all populations were not significant (*p*=0.05; Table 4; Fig 5). In addition, negative Fu’s F_*S*_ estimates were observed in pigs in the Philippines (−1.503; *p*=0.05), Laos (−2.73; *p*=0.05) and Vietnam (−0.0499; *p*=0.05). The Ramos-Onsins & Rozas’ R^2^ tests [39] were not significant in all cases, except for the MSEA samples combined (*p*<0.05). The null hypothesis of expectation under the sudden expansion model such as the sum square deviation and raggedness test result (*p*=0.05) on coalescent estimates, was not statistically supported in all subject locations, except for the combined MSEA samples. The analysis of the prehistoric population size dynamics of Philippine native pigs using BSP was consistent with the result from the mismatch distribution analyses. As projected by the BSP, the Philippine native pig population revealed a long stationary period of effective population size (Fig 6A). The sudden population decrease event occurred roughly at about ∼25,000 years before present (BP). At the regional scale, MSEA pigs marked a gradual significant increase in population approximately during the Late Pleistocene ages (Fig 6B).

**Fig 4.**
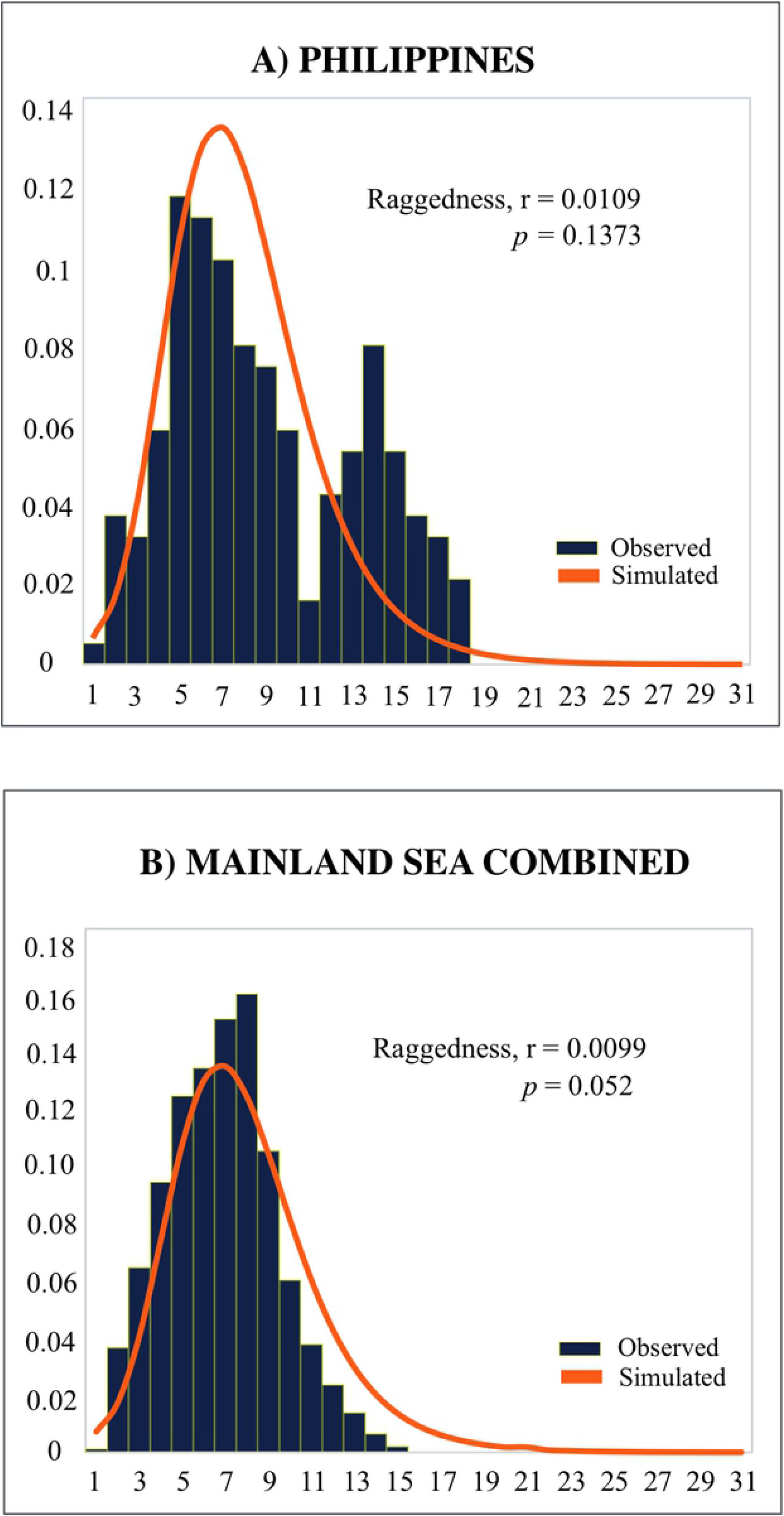
**Mismatch distributions of mitochondrial DNA sequences of the (A) Philippine pigs, (B) mainland SEA pigs based on pairwise nucleotide differences.**

**Table 4.** Values of neutrality test (Fu’s F_*S*_ and Tajima’s D), sum of square deviation (SSD) and Harpending’s raggedness index (H*ri*) for Philippine native pigs and MSEA pig mtDNA D-loop region

| COUNTRIES | Fu's $F_S$ test | Tajima's $D$ test | SSD | $H_{ri}$ |
| --- | --- | --- | --- | --- |
| KHM | 0.421 | 1.400 | 0.054* | 0.101* |
| BTN | 1.049 | 0.678 | 0.048 | 0.089 |
| LAO | -2.73 | 0.264 | 0.035 | 0.061 |
| MMR | -6.542* | -0.061 | 0.004 | 0.016 |
| VNM | -0.499 | 0.768 | 0.037 | 0.129 |
| TOTAL | -25.282** | -0.014 | 0.006 | 0.012 |
| PHL | -1.503 | 0.231 | 0.017 | 0.020 |

\* $p < 0.05$ ; \*\* $p < 0.01$ as tested by randomization (1000 permutations) using ARLEQUIN. PHL= Philippines; KHM= Cambodia; BTN= Bhutan; MMR= Myanmar; VNM= Vietnam.

**Fig 5.**
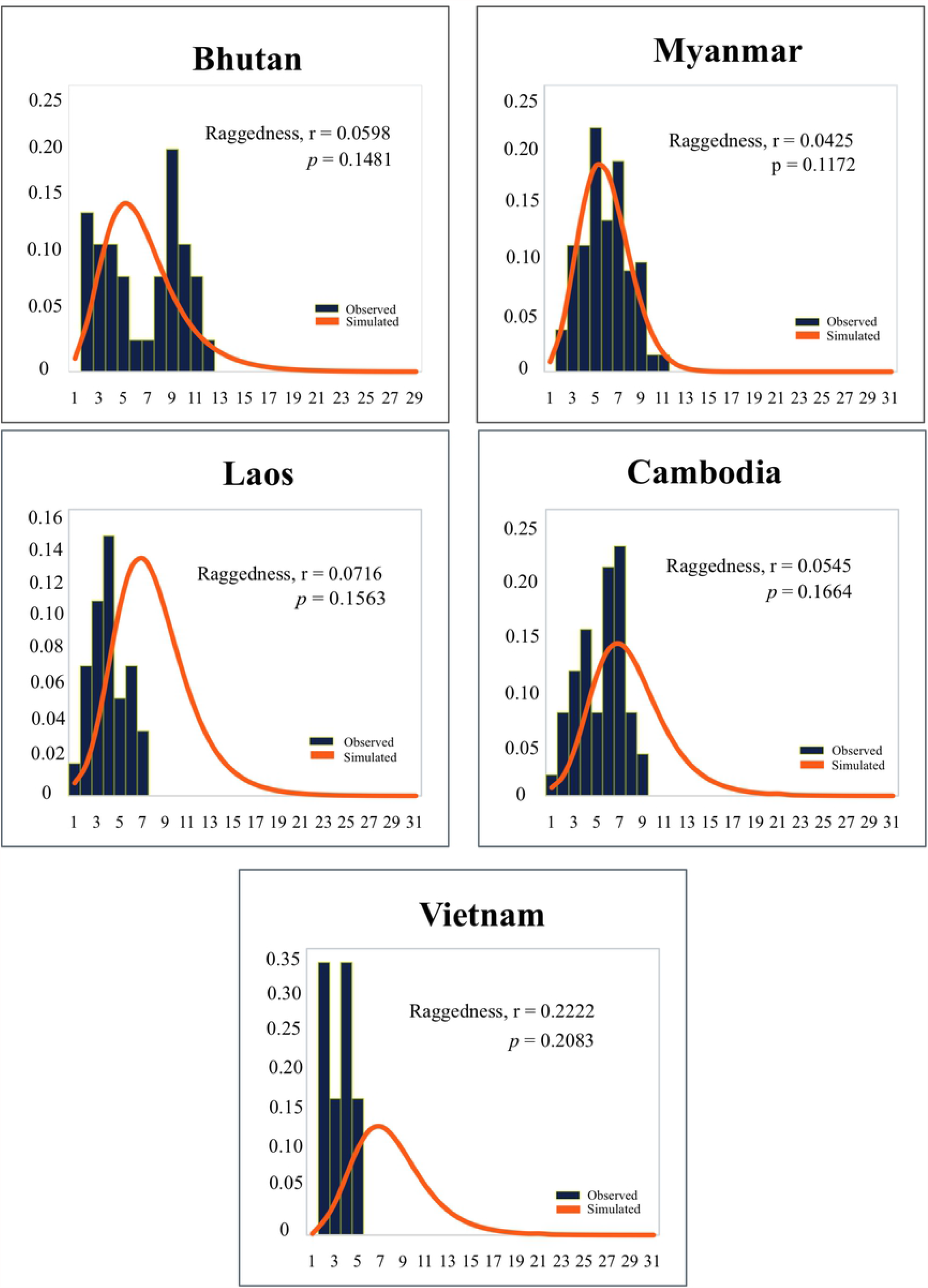
**Mismatch distributions of mitochondrial DNA sequences of countries in mainland Southeast Asian pigs based on pairwise nucleotide differences.**

**Fig 6.**
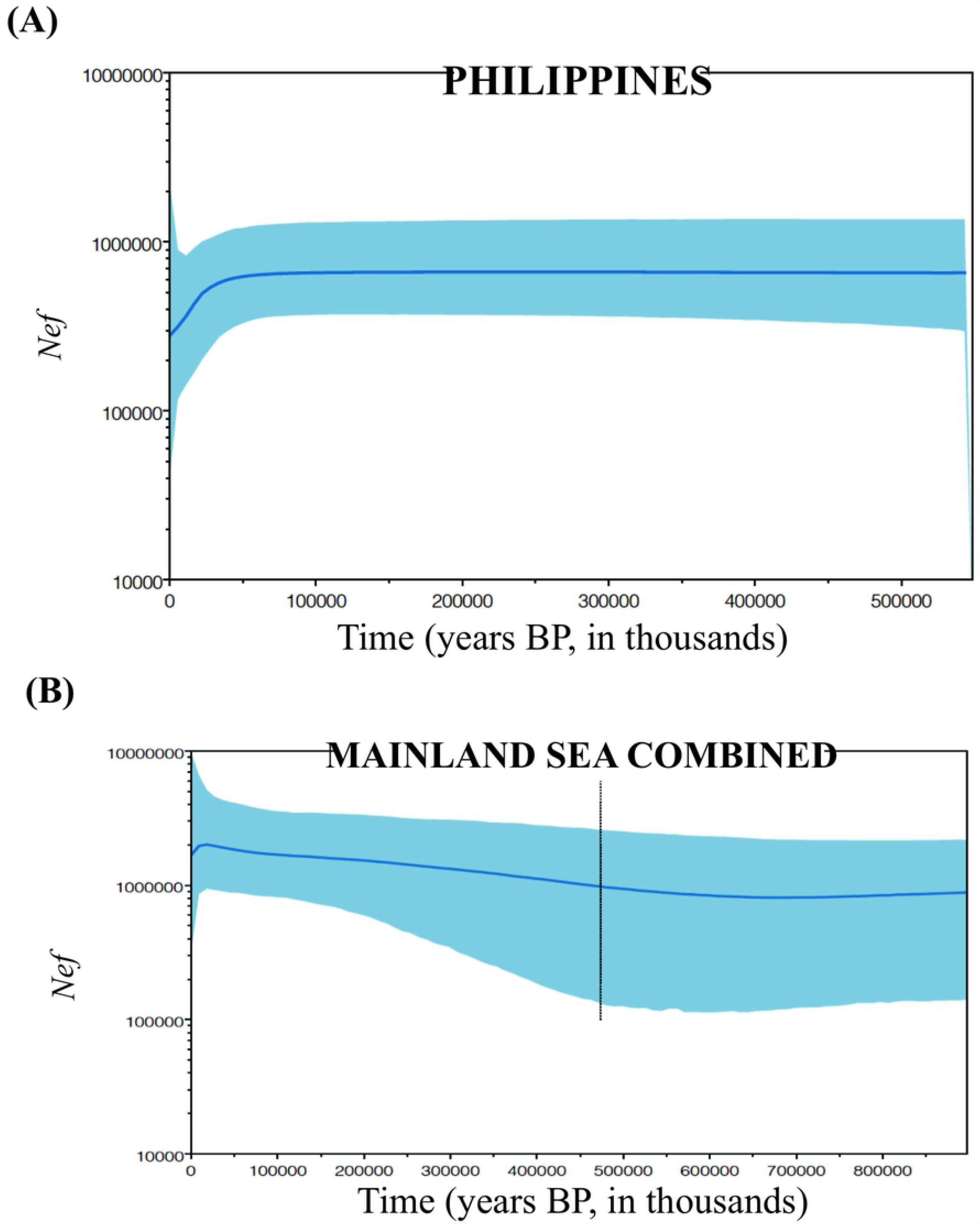
**Bayesian skyline plots showing effective population size of (A) Philippine and (B) mainland Southeast Asian pigs. Median estimates of female effective population size (*Nef*) are shown as solid thick line (blue) and the light-blue shaded area marks the 95% credibility intervals. The abscissa is scaled in thousands of years before present (BP). The Philippine pigs revealed a long stationary period of effective population size and the sudden population decrease event occurred roughly at about ∼25**,**000 BP.**

## Discussion

Literature on genetic studies in Philippine pigs is scarce, although this animal represents excellent genetic resources for the local economy and underlies as a genetic basis to study human settlements and migration. This study provides the first comprehensive data on the history of dispersal, genetic structure and diversity, and population dynamics of Philippine native pigs. Previous studies have revealed multiple centers of pig domestication (six major clusters, denoted as D1 to D6) and the existence of a clear phylogenetic structure of mtDNA D-loop region sequences found in wild boars and domestic pigs [3-4], and their possible association with the hypothesized Neolithic expansion in Island South East Asia and Oceania [4]. Therefore, the patterns of haplotype distribution seen in this study can be associated with the importance of identifying the prehistoric arrival of domestic pigs in the Philippines, and it’s spread across the islands as rooted to the hypothesized migratory route of Neolithic Austronesian-speaking populations from Taiwan into the Philippines [47-48] and the possible pig dispersal that occurred from ISEA via Palawan and the Sulu Archipelago. The phylogenetic patterns in the current study generally agreed on the existing two core lineages, one of the Asian phylogeographic origins and one of the European phylogeographic origins. The distribution of the haplotype frequencies indicated no equilibrium thus, the geographically distributed haplotypes suggested that the present-day Philippine native pigs have multiple ancestral origins spread across the Eurasian Continent. The close genetic connection between the continental wild boars and domestic pigs from the MSEA and NEA present in the Philippine pig genetic pool corroborates our hypothesis of a genetic signal that could potentially be associated with the recently reported multiple waves of human migrations to the Philippines during the last 50,000 years [49]. During the glacial periods, extensive gene flow has suggested to occurred among the *Sus* species [50] and was considered as an important driving factor that has established the present-day geographic distribution of *Sus* populations throughout the world [51]. Thus, these events may have paved the way for these pigs with multiple ancestral lineages to be introduced in the country, including domestic animals like chickens [52], goats [53], cattle [54], and other species that have adapted to local conditions and developed distinctive characteristics. While the preliminary studies that revealed the absence of the Pacific Clade in the Philippine Archipelago [24] have led some researchers to challenge the veracity of pig dispersal from the NEA throughout the Pacific Islands via the Philippines [22]. This study reported the first Pacific Clade signature and the ubiquitously distributed D2 haplotypes which could potentially shed light on the question of pig dispersal by the Austronesian-speaking populations from NEA via the Philippines. Furthermore, the close genetic association of three haplotypes (PHL15,22-23) to Lanyu pigs of Taiwan strongly suggests an mtDNA maternally derived from the same lineage. Therefore, these patterns of the genetic variation in contemporary Philippine pigs could mirror the multilayered history of the Philippines as a nation with a rich history of trade and bartering of voyagers with coastal communities, including the riverine movements into near-coastal settlements during prehistoric and protohistoric times [55] that has contributed significantly to the genetic landscape of the Asia-Pacific region [49].

Before the European arrived in the Philippines during the Spanish colonization, there was indication that pigs were introduced already by the Chinese traders [56], and subsequently followed by the intensive importation of various exotic pig breeds from Europe [57] that has resulted in a diversified Philippine pig genetic pool. This hypothesis is precisely evident as shown by the close genetic relationships between Philippine native pigs and Chinese pigs, which exhibited by the similarities in their morphology and mtDNA variation due to introgression. It is undeniable that the Chinese mtDNA footprint was imperative in the history of Philippine native pig development. Compared to the high signal of genetic introgression of Chinese pig breeds into Philippine native pigs, European pig maternal introgression was minimal which constitute only about 5.10% of the studied population. This observation was congruent with the situation among domestic pigs in MSEA where the mtDNA of European pigs was considered to have a negligible impact on the maternal origin of domestic and native pigs. Considering our sampling as aggregates of both lowland and upland areas, our visual observation and molecular result implies that the exotic pig breeds have not yet fully penetrated the remote areas in the Philippines. While the pervasive indiscriminate hybridization between exotic pig breeds and native pigs (i.e., Berkshire or Duroc x Philippine native pigs) in the lowland areas could pose an important challenge from a long-term management perspective. Previously, it was underscored that the European pigs have both Asian and European pig mtDNA, resulted from the extensive history of crossbreeding between European and Asian pigs with the predominance of Asian mtDNA introgression [58-59,3] and now constitutes about 20-35% of Asian matrilineal origin [60-62]. Hence, this concurrent maternal introgression of the global breeds, which currently contributes about 30% in D2 haplogroup and widely distributed in Chinese pigs, cannot be rejected, as it was evident in the maternal signatures of domestic pigs in almost all of Asia.

Our results indicated a high proportion of Philippine pig haplotypes (49.07%; 53/108 individuals) that fell under the D7 haplogroup compared to the previously identified similar haplotypes in MSEA (34.62%; 45/130 individuals). This haplogroup was not previously reported in Chinese pigs and has precluded the origin of these haplotypes out of China [44]. As reported earlier, this haplogroup was the most recent pig mtDNA lineage discovered, distinct from those in previously documented centers of pig domestication. Due to the absence of a similar haplotype in the Insular and NEA regions, the phylogeographic origin of the Phil-D7 (Philippine type-D7 haplotypes) represents an interesting question, as its haplotype distribution does not primarily suggest the IBBH as the direct origin of the Phil-D7 or the probability of whether these haplotypes were established as a result of a human-mediated introduction into the Philippines. Further, the pattern of the distribution of haplotypes does not coincide within the hypothesized migratory route of the Neolithic Austronesian-speaking populations, and likewise with the possible pig ancestral diffusion that occurred from the ISEA to the Philippines sometime during the interglacial periods of the Pleistocene. While recent studies on combining mitochondrial DNA and geometric morphometric of the pig have shown some human-mediated dispersal in some islands of Southeast Asia [63,16,23,25], there is presently no genetic data nor archaeological material that provide evidence for any prehistoric translocation of Philippine pigs between the islands of the archipelago [24,64]. However, assuming that haplotypes between these two geographic locations originate from one ancestral lineage, the significant population differentiation, despite sharing of haplotypes, could be suspected as a consequence of geographical isolation. The vicariance brought about by the severe Quaternary Sea level drops resulted in the isolation of populations due to the formation of geographic barriers to migration and consequent genetic divergence between these populations [64]. Population differentiation usually occurs when there is migration of a certain population away from its founder population which leads to a reduction in genetic diversity as predicted by the theory of Genetic Isolation by Distance. Moreover, an expansion model from a single founder predicts that patterns of genetic diversity in populations can be thoroughly explained by their geographic expansion from the founders, concomitant to genetic differentiation [65].

The Philippine wild pigs are known or reported from all of the larger and many of the smaller, offshore islands in the Philippines [15]. In this study, the relatively high degree of genetic variation detected in Philippine wild pig haplotypes between the observed populations including samples from the GenBank is compelling. The phylogenetic tree and haplotype network analyses illustrated an extremely vague cluster among other wild boars. Therefore, in this current study, it seems that the mtDNA of Philippine wild pig does not have a significant maternal contribution to Philippines native pigs disparate from the limited information suggesting that it was derived from the numerous wild pigs in the country.

The haplotype diversity of the studied Philippine native pig population was generally moderate and similar with those in the MSEA countries such as Laos and Myanmar, while relatively higher compared to those from Cambodia, Bhutan, and Vietnam. The nucleotide diversity, however, was remarkably higher in Philippine native pigs. As emphasized previously, nucleotide diversity represents a more suitable parameter than haplotype diversity in estimating the genetic diversity in a population [66], as it addresses both the frequency of haplotypes and the nucleotide differences between haplotypes. These values were relatively higher in the previously reported pig nucleotide diversity in southern China including Yunnan, the Tibetan highlands, the extensive basins of the Yangtze and Yellow Rivers, Taiwan and some Pacific Islands. These are similar to those found in the outlying areas of ISEA, Korea [67,62] and Bhutan. This pattern of genetic variation suggests a scenario that reflects past expansion dynamics from the species area of origin [68], owing to subsequent translocations by humans and the effects of introgression between different DNA lineages [69]. Also, [68] have previously discussed that the present large-scale pattern of genetic variability in *Sus* can be linked to one or more ancient long-distance colonization events followed by divergence of isolated lineages, geographical extinction due to local extinction within a previously continuous distributional range, and isolation by distance which resulted in restricted gene flow.

The genetic fixation observed in the studied Philippine native pig populations, reported as F_*ST*_, indicated that gene flow is limited among individuals which suggested populations between regions are genetically isolated from each other. Nevertheless, this is an expected population scenario for the Philippine native pigs where the natural genetic exchange is confined as a result of the archipelagic geographical setting of the Philippines. During the past glacial periods, the Philippine archipelago was believed to have never been connected to the Asian continent [15-16,14], which has influenced restricted genetic exchange and mtDNA distribution of pigs throughout the islands.

### Past population dynamics

The historical demography of Philippine native pig populations was examined using mismatch distributions which represent the frequency distribution of pairwise differences among all sampled haplotypes. Theoretical studies have shown that population bottlenecks and population expansions cause a sound effect on the pattern of genetic polymorphism among haplotypes in the population [35]. The multimodal pattern of the mismatch distribution in Philippine native pigs can be assumed to have undergone irrelevant demographic expansion that have occurred over a long time and that the population could have been shown long-term stability or decline. In addition, the neutrality test based on both the Tajima’s D (*p*=0.05) and Fu’s F_*S*_ statistics (*p*=0.05) was not consistent with the recent population and demographic expansions. The Fu’s F_*S*_ test is highly sensitive to demographic expansion which results in large negative F_*S*_ values, whereas the significant Tajima’s D value could be a sign of population expansion and bottleneck [70-72]. The R^2^ statistics and the simulation based on coalescent process as quantified by the raggedness index confirmed the mismatch distribution of the studied populations. The haplotypic and genealogical relationship portrayed in the reduced-median network, despite a significant population subdivision among population, also showed no geographic structuring except for a few star-like patterns. In general, a population that has gone through a recent population expansion displays a star-like structure in a network tree, smooth, and unimodal mismatch distribution [38] because most alleles descended from one or a few ancestral types [35]. Therefore, the reflected mismatch distribution could be a signature of the presence of different haplogroups detected rather than a demographic stability.

The Bayesian skyline plot revealed a sudden decrease in population during the interglacial periods of the Late Pleistocene. The cyclic sea level fluctuations during these periods are regarded as one of the most important events involved in shaping the contemporary geographic distribution of genetic variation and evolutionary dynamics of the population [73-75]. Moreover, the alleged decline in the population of some animals has occurred because of glacial-interglacial episodes [76-77]. This is a consistent population scenario experienced by both Asian and European wild boars, where they experienced population bottlenecks during the Last Glacial Maximum (LGM; ∼20,000 years ago). A considerable drop in population size was more pronounced in Europe than in Asia which has caused the low genetic diversity seen in modern European wild boars [60]. The recent Ice Age has also caused a huge sea level drop of about 120 m below the present level, exposing huge areas as dry land, but the Philippines remained isolated by deep channels. Conceivably, this influenced the land distribution of pigs in the Philippine archipelago that resulted to their geographic isolation and subsequently restricted gene flow. The effects of bottlenecks are evident in populations occupying smaller geographic ranges which are vulnerable to stochastic events and genetic drift compared to larger and more widespread populations [78]. Our result of a population expansion in MSEA pig was in contrary with the population scenario reported by [50], and a more severe bottleneck in ISEA during the Pleistocene periods. These population declines are consistent with the reduction of temperature during this period that would have reduced the overall forest cover in these areas [79-80,50].

## Conclusion

This study provided critical insights that will properly help address the contradicting hypothesis of a possible human-mediated translocation and exchange of pigs involving the Philippines. The results of our study could support the Neolithic-Austronesian model of expansion while a more rigorous investigation should be carried out in linking the possible pig ancestral diffusion that took place from MSEA via Sundaland to the Philippines. The unique geographical features of the Philippines have resulted in an insignificant migration flow of pigs and as a long-term consequence, geographical isolation had occurred. The underlying sudden population decline as predicted in the BSP markedly followed the LGM period. Ultimately, the escalating rate of hybridization of Philippine native pigs with commercial stocks in the Philippines represents a severe risk for native pig populations. Therefore, urgent conservation measures and suitable management of their genetic pool are crucial in the management of animal genetic resources at the local and global levels. For future perspectives, Y-specific markers could be performed to assess the level of male-mediated introgression of European pigs into Philippine native pigs.

## Acknowledgements

We are indebted to Dr. Lawrence M. Liao for the technical support and numerous insights that greatly improved this work. We thank Cyrill John Prima Godinez, Jant Cres Caigoy, and all the members of the Animal Genetics Laboratory of Hiroshima University. We thank the Visayas State University, Capiz State University, Aklan State University, Iloilo State College of Fisheries, farmers, and backyard pig raisers throughout the Visayas for the help during our sampling.

## Supporting information

**S1 Fig. Variable positions among haplotypes of the partial mitochondrial DNA control region (about 510 bp) found in this study**. Dots (.) indicates matches with the nucleotide sequence GenBank accession number AB041480 (Main cluster of Asian origin); minus (−) represents gaps. PWP=Philippine wild pigs; D7=previously described as MTSEA haplogroup. Nucleotide positions are numbered according to our sequence alignment.

**S2 Fig. Proposed route of dispersal and human-mediated translocation of pigs in the Philippines**.

**S1 Table. List of samples used in the study**.

**S2 Table. Newly generated Philippine pig haplotypes and the publicly available sequences of pigs found in the mainland Southeast Asia**

**S3 Table. Publicly available global pig haplotype sequences used to infer phylogenetic and network haplotypes analysis**

## References

1. Ruvinsky A, Rothschild MF, Larson G, Gongora J. Systematics and evolution of the pig. In: Rothschild MF, Ruvinsky A, (eds). The Genetics of the Pig. Oxforshire: CAB International; 2011. pp. 1–13.

2. Epstein J, Bichard M. Pig In: Evolution of Domesticated Animals. New York, NY: Longman; 1984. pp. 145–162.

3. Giuffra E, Kijas JMH, Amarger V, Calborg O, Jeon J-T, Andersson L. The origin of the domestic pig: Independent domestication and subsequent introgression. Genetics. 2000;154: 1785–1991.

4. Larson G, Dobney K, Albarella U, Fang M, Smith EM, Robins J, Lowden S, Finlayson H, Brand T, Willerslev E, Conwy PR, Andersson L, Cooper A. Worldwide phylogeography of wild boar reveals multiple centers of pig domestication. Science. 2005;307: 1618–1621.

5. Scandura M, Iacolina L, Crestanello B, Pecchioli E, Di Benedetto MF, Russo V, Davoli R, Apollonio M, Bertorelle G. Ancient vs. recent processes as factors shaping the genetic variation of the European wild boar: Are the effects of the last glaciation still detectable? Molecular Ecology. 2008;17: 1745–1762.

6. Vilaca ST, Biosa D, Zachos F, Iacolina L, Kirschning J, Alves PC. Mitochondrial phylogeography of the European wild boar: The effect of climate on genetic diversity and spatial lineage sorting across Europe. Journal of Biogeography. 2014;41(5): 987– 998.

7. Frantz LAF, Meijaard E, Gongora J, Haile J, Groenen MAM, Larson G. The evolution of Suidae. Annual Review of Animal Bioscience. 2016;4(1): 61–85.

8. Ramirez O, Ojeda A, Tomàs A, Gallardo D, Huang LS, Folch JM. Integrating Y-chromosome, mitochondrial, and autosomal data to analyze the origin of pig breeds. Molecular Biology and Evolution. 2009;26(9): 2061–2072.

9. Amills M, Megens H-J, Manunza A, Ramos-Onsins SE, Groenen MA. A genomic perspective on wild boar demography and evolution. In: Meletti M, Meijaard E, editors. Ecology, Conservation and Management of Wild Pigs and Peccaries. Cambridge University Press; 2017. pp. 376–387.

10. Heaney L, Mittermeier RA. 1997. The Philippines. In: Mittermeier RA, Mittermeier CG, Robles GP. Me (eds) Megadiversity: Earth’s Biologically Wealthiest Nations. Monterrey (Mexico), CEMEX; 1997. pp. 236–255.

11. Myers N, Mittermeier RA, Mittermeier CG, da Fonseca Gab, Kent J. Biodiversity hotspots for conservation priorities. Nature. 2000;403: 853–858.

12. Roberts CM. Marine biodiversity hotspots and conservation priorities for tropical reefs. Science. 2002;295: 1280–1284.

13. Posa MRC, Diesmos AC, Sodhi NC, Brooks TM. Hope for threatened tropical biodiversity: Lessons from the Philippines. Bioscience. 2008;58(3): 231–240.

14. Voris HK. Maps of Pleistocene sea levels in Southeast Asia: Shorelines, river systems and time durations. Journal of Biogeography. 2000;27: 1153–1167.

15. Oliver WLR. The taxonomy, distribution and status of Philippine wild pigs. Ibex, Journal of Mountain Ecology. 1995;3: 26–32.

16. Groves CP. Taxonomy of wild pigs (Sus) of the Philippines. Zoological Journal of the Linnaean Society. 1997;120: 163–191.

17. Li KY, Li KT, Yang CH, Hwang MH, Chang SW, Lin SM, Wu HJ, Basilio EB, Vega RSA, Laude RP, Ju YT. Insular East Asia pig dispersal and vicariance inferred from Asian wild boar genetic evidence. Journal of Animal Science. 2017;95(4): 1451–1466.

18. Bellwood P and Dizon E. The Batanes Archaeological Project and the ‘Out of Taiwan’ hypothesis for Austronesian dispersal. Journal of Austronesian Studies. 2005;1(1): 1– 32.

19. Bellwood P and Dizon E. The chronology of Batanes prehistory. In P. Bellwood, and E. Dizon (eds), 4000 Years of Migration and Cultural Exchange: Archaeology in the Batanes Islands, Northern Philippines. Terra Australis 40. Canberra: ANU E Press; 2013. pp. 67–76.

20. Oppenheimer S. Eden in the East. Phoenix: Orion; 1999.

21. Solheim WG. Archaeology and culture in Southeast Asia: unravelling the Nusantau. Quezon City: University of the Philippines Press; 2006.

22. Larson G, Albarella U, Dobney K, Rowley-conwy P. Current views on Sus phylogeography and pig domestication as seen through modern mtDNA studies, in U. Alberella, K. Dobney, A. Ervynck & P. Rowley-conwy (eds.) Pigs and humans: 10,000 years of interaction. Oxford: Oxford University Press; 2007b. pp. 30–41.

23. Larson G, Cucchi T, Fujita M, Smith EM, Robins J, Anderson A, Rolett B, Spriggs M, Dolman G, Kim TH, Thuy NTD, Randi E, Doherty M, Due RA, Bollt R, Djubiantono T, Griffin B, Intoh M, Keane E, Kirch P, Li KT, Morwood M, Pedrina LM, Piper PJ, Rabell RJ, Shooter P, den Bergh GV, West E, Wickler S, Yuan J, Cooper A, Dobney K. Phylogeny and ancient DNA of Sus provides insights into Neolithic expansion in island Southeast Asia and Oceania. Proceedings of the National Academy of Sciences. 2007a;104: 4834–4839.

24. Dobney K, Cucchi T, Larson G. The pigs of Island Southeast Asia and the Pacific: New evidence for taxonomic status and human-mediated dispersal. Asian Perspectives. 2008;47(1): 59–74.

25. Larson G, Liu R, Zhao X, Yuan J, Fuller D, Barton L, Dobney K, Fan Q, Gu Z, Liu XH, Luo Y, Lv P, Andersson L, and Li N. Patterns of East Asian pig domestication, migration, and turnover revealed by modern and ancient DNA. Proceedings of the National Academy of Sciences. 2010;107: 7686–7691.

26. Nishibori M, Hanazono M, Yamamoto Y, Tsudzuki M, Yasue H. Complete nucleotide sequence of mitochondrial DNA in White Leghorn and White Plymouth Rock chickens. Animal Science Journal. 2003;74(5): 437–439.

27. Nishibori M, Hayashi T, Tsudzuki M, Yamamoto Y, Yasue H. Complete sequence of the Japanese quail (Coturnix japonica) mitochondrial genome and its genetic relationship with related species. Animal Genetics. 2001;32(6): 380–385.

28. Tamura K, Stecher G, Peterson D, Filipski A, Kumar S. MEGA6: Molecular evolutionary genetics analysis version 6.0. Molecular Biology and Evolution. 2013;30(12): 2725–2729.

29. Librado P, Rozas J. A software for comprehensive analysis for DNA polymorphism data. Bioinformatics. 2009;25: 1450–1452.

30. Excoffier L, Laval G, Schneider S. Arlequin ver. 3.0. An integrated software package for population genetics data analysis. Evolutionary Bioinformatics Online. 2005;1: 47– 50.

31. Excoffier L, Smouse P, Quattro J. Analysis of molecular variance inferred from metric distances among DNA haplotypes: Application to human mitochondrial DNA restriction data. Genetics. 1992;131: 479–491.

32. Guindon S, Dufayard JF, Lefort V, Anisimova M, Hordijk W, Gascuel O. New algorithm and methods to estimate Maximum-Likelihood Phylogenies: Assessing the permormance of PhyML 3.0. Systematic Biology. 2010;59(3): 307–321.

33. Bandelt HJ, Forster P, Sykes BC, Richards MB. Mitochondrial portraits of human populations. Genetics. 1995;141: 743–753.

34. Bandelt HJ, Forster P, Rolh A. Median-joining networks for inferring intraspecific phylogenies. Molecular Biology and Evolution. 1999;16: 37–48.

35. Rogers AR, Harpending H. Population growth makes waves in the distribution of pairwise genetic differences. Molecular Biology and Evolution. 1992;9: 552–569.

36. Harpending HC. Signature of ancient population growth in a low-resolution mitochondrial DNA mismatch distribution. Human Biology. 1994;66: 591–600.

37. Rogers AR. Genetic evidence for a Pleistocene population explosion. Evolution. 1995;49: 608–615.

38. Fu YX. Statistical tests of neutrality of mutations against population growth, hitchhiking and background selection. Genetics. 1997;147(2): 915–925.

39. Ramos-Onsins SE, Rozas J. Statistical properties of the new neutrality test against population growth. Molecular Biology and Evolution. 2002;19: 2092–2100.

40. Drummond AJ, Rambaut A, Shapiro B, Pybus OG. Bayesian coalescent inference of past population dynamics from molecular sequences. Molecular Biology and Evolution. 2005. 22: 1185–1192.

41. Hasegawa M, Kishino H, Yano T. Dating of the human-ape splitting by a molecular clock of mitochondrial DNA. Journal of Molecular Evolution. 1985;22: 160–174.

42. Bouckaert R, Heled J, Kühnert D, Vaughan T, Wu CH, Xie D, Suchard MA, Rambaut A, Drummond AJ. BEAST 2: A software platform for Bayesian evolutionary analysis. PLoS Computational Biology. 2014;10: e1003537.

43. Pesole G, Gissi C, De Chirico A, Saccone C. Nucleotide substitution rate of mammalian mitochondrial genomes. Journal of Molecular Evolution. 1999;48: 427–433.

44. Tanaka K, Iwaki Y, Takizawa T, Dorji T, Tshering G, Kurosawa Y, Maeda H, Mannen H, Nomura K, Dang VB, Chhum-Phith LC, Bouahom B, Yamamoto Y, Dang T, Namikawa T. Mitochondrial diversity of native pigs in the mainland South and South-east Asian countries and its relationships between local wild boars. Animal Science Journal. 2008;79: 417–434.

45. Okumura N, Kurosawa Y, Kobayashi E,Watanabe T, Ishiguro N, Yasue H, Mitsuhashi T. Genetic relationship amongst the major non-cording regions of mitochondrial DNAs in wild boars and several breeds of domesticated pigs. Animal Genetics. 2001;32: 139– 147.

46. Fang M, Andersson L. Mitochondrial diversity in European and Chinese pigs is consistent with population expansions that occurred prior to domestication. Proceedings Biological Sciences of the Royal Society. 2006;273: 1803–1810.

47. Bellwood P. A hypothesis for Austronesian origins. Asian Perspectives. 1985;26: 107– 118.

48. Ingicco T, Piper PJ, Amano N, Paz VJ, Pawlik AF. Biometric differentiation of wild Philippine pigs from introduced Sus scrofa in modern and archaeological assemblages. International Journal of Osteoarchaeology. 2017;27: 768–784.

49. Larena M, Quinto FS, Sjödin P, McKenna J, Ebeo C, Reyes R, Casel O, Huang J-Y, Hagada KP, Guilay D, Reyes J, Allian FP, Mori V, Azarcon LH, Manera A, Terando C, Jamero L, Sireg G, Manginsay-Tremedal R, Labos MS, Vilar RH, Latiph A, Saway RS, Marte E, Magbanua P, Morales A, Java I, Reveche R, Barrios B, Burton E, Salon JC, Kels MJT, Albano A, Cruz-Angeles RB, Molanida E, Granehäll L, Vicente M, Edlund H, Loo JH, Ho TYW, Reid L, Malmström H, Schlebusch C, Lambeck K, Endicott P, Jakobsson M. Multiple migrations to the Philippines during the last 50,000 years. Proceedings of the National Academy of Sciences, USA. 2021;118(13): e2026132118.

50. Frantz LAF, Schraiber JG, Madsen O, Megens HJ, Bosse M, Paudel Y, et al. Genome sequencing reveals fine scale diversification and reticulation history during speciation in Sus. Genome Biology. 2013;14(9): R107.

51. Gongora J, Fleming P, Spencer PBS, Mason R, Garkavenko O, Meyer JN, Droegemueller C, Lee JH, Moran C: Phylogenetic relationships of australian and new zealand feral pigs assessed by mitochondrial control region sequence and nuclear gpip genotype. Molecular Phylogenetics and Evolution. 2004;33: 339–348.

52. Thomson VA, Lebrasseur O, Austin JJ, Hunt TL, Burney DA, Denham T, Rawlence NJ, Wood JR, Gongora J, Girdland Flink L, Linderholm A, Dobney K, Larson G, Cooper A. Using ancient DNA to study the origins and dispersal of ancestral Polynesian chickens across the Pacific. Proceedings of the National Academy of Sciences, USA. 2014;1: 111(13):4826–31.

53. Naderi S, Rezaei H-R, Taberlet P, Zundel S, Rafat S-A, Naghash H-R, Elbarody MAA, Ertugrul O, Pompanon F, for the Econogene Consortium. Large-scale mitochondrial DNA analysis of the domestic goat reveals six haplogroups with high diversity. PLoS ONE. 2007;2(10): e1012.

54. Scott WH. Sixteenth-century Visayan food and farming. Philippine Quarterly of Culture and Society. 1990;18: 291–293.

55. Fox RB. The Archeological record of Chinese influences in the Philippines. Philippine Studies. 1967;15(1):41–62.

56. Penalba FF. Philippine native pigs: Performance and potential. Animal Production Technology. 1993;8(1): 2–7.

57. Bondoc OL. Animal Breeding: Principles and Practice in the Philippine Context. Diliman, Quezon City: University of the Philippines Press; 2008.

58. Watanabe T, Hayashi Y, Ogasawara N, Tomoita T. Polymorphism of mitochondrial-DNA in pigs based on restriction endonuclease cleavage patterns. Biochemical Genetics. 1985;23: 105–113.

59. Okumura N, Ishiguro N, Nakano M, Hirai K, Matsui A, Sahara N. Geographic population structure and sequence divergence in the mitochondrial DNA control region of the Japanese wild boar (Sus scrofa leucomystax), with reference to those of domestic pigs. Biochemical Genetics. 1996;34: 179–189.

60. Groenen M, Archibald A, Uenishi H, et al. Analyses of pig genomes provide insight into porcine demography and evolution. Nature. 2012;491: 393–398.

61. Bianco E, Soto HW, Vargas L, Pérez-Enciso M. The chimerical genome of Isla del Coco feral pigs (Costa Rica), an isolated population since 1793 but with remarkable levels of diversity. Molecular Ecology. 2015;24: 2364–2378.

62. Wang C, Chen YS, Han JL, M., Li XJ, Liu X-H. Mitochondrial DNA diversity and origin of indigenous pigs in South China and their contribution to western modern pig breeds. Journal of Integrative Agriculture. 2019;18(10): 2338–2350.

63. Groves CP. Pigs east of the Wallace Line. Journal de la Soci’et’e des Oc’eanistes. 1983;39: 105–119.

64. Baker PA, Fritz SC, Dick CW, Eckert AJ, Horton BK, Manzoni S, Ribas CC, Garzione CN, Battisti, DS. The emerging field of geogenomics: Constraining geological problems with genetic data. Earth-Science Reviews. 2014;135: 38–47.

65. Dorcus KM, Steffen W, Armin OS, Annett W, Christian R, Henner S. Genetic diversity in global chicken breeds as a function of genetic distance to the wild populations. 2020. Preprint; https://doi.org/10.1101/2020.01.29.924696.

66. Sulandari S, Zein MA, Sartika T. Molecular characterization of Indonesian indigenous chickens based on mitochondrial DNA displacement (D) -loop sequences. Hayati Journal of Bioscience. 2008;15: 145–154.

67. Wu GS, Yao YG, Qu KX, Ding ZL, Li H, Palanichamy MG, Duan ZY, Chen YS, Zhang YP. Population phylogenomic analysis of mitochondrial DNA in wild boars and domestic pigs revealed multiple domestication events in East Asia. Genome Biology. 2007b;8: R245.

68. Scandura M, Iacolina L, Apollonio M. Genetic diversity in the European wild boar Sus scrofa: Phylogeography, population structure and wild x domestic hybridization. Mammal Review. 2011;41(2): 125–137.

69. Bruford M, Bradley D, Luikart G. DNA markers reveal the complexity of livestock domestication. Nature Reviews Genetics. 2003;4: 900–910.

70. Tajima F. The effect of change in population size on DNA polymorphism. Genetics. 1989;123: 597–601.

71. Fauvelot C, Bernardi G, Planes S. Reductions in the mitochondrial DNA diversity of coral reef fish provide evidence of population bottlenecks resulting from Holocene sea-level change. Evolution. 2003;57: 1571–1583.

72. Martel C, Viard F, Bourguet D, Garcia-Meunier P. Invasion by the marine gastropod Ocinebrellus inornatus in France. II. Expansion along the Atlantic coast. Marine Ecology Progress Series. 2004;273: 163–172.

73. Hewitt GM. The genetic legacy of the quaternary ice ages. Nature. 2000;405: 907–913.

74. Provan J, Bennett KD. Phylogeographic insights into cryptic glacial refugia. Trends in Ecology and Evolution. 2008;23: 564–571.

75. Deli T, Kiel C, Schubart CD. Phylogeographic and evolutionary history analyses of the warty crab Eriphia verrucosa (Decapoda, Brachyura, Eriphiidae) unveil genetic imprints of a late Pleistocene vicariant event across the Gibraltar Strait, erased by postglacial expansion and admixture among refugial lineages. BMC Evolutionary Biology. 2019;19: 105.

76. Lyrholm T, Leimar O, Gyllensten U. Low diversity and biased substitution patterns in the mitochondrial DNA control region of sperm whales: Implications for estimates of time since common ancestry. Molecular Biology and Evolution. 1996;13: 1318–1326.

77. Sanchez G, Kawai K, Yamashiro C, Fujita R, Wakabayashi T, Sakai M, Umino T. Patterns of mitochondrial and microsatellite DNA markers describe historical and contemporary dynamics of the Humboldt squid Dosidiscus gigas in the Eastern Pacific. Reviews in Fish Biology and Fisheries. 2020;30: 519–533.

78. Velickovic N, Ferreira E, Djan M. Demographic history, current expansion and future management challenges of wild boar populations in the Balkans and Europe. Heredity. 2016;117: 348–357.

79. Bird MI, Taylor D, Hunt C. Palaeoenvironments of insular Southeast Asia during the Last Glacial Period: a savanna corridor in Sundaland? Quaternary Science Review. 2005;24: 2228–2242.

80. Wurster CM, Bird MI, Bull ID, Creed F, Bryant C, Dungait JAJ, Paz V. Forest contraction in north equatorial Southeast Asia during the Last Glacial Period. Proceedings of the National Academy of Sciences. 2010;107: 15508–15511.

